# Physiological properties and tailored feeds to support aquaculture of marbled crayfish in closed systems

**DOI:** 10.1101/2020.02.25.964114

**Authors:** Sina Tönges, Karthik Masagounder, Julian Gutekunst, Jasmin Lohbeck, Aubry K. Miller, Florian Böhl, Frank Lyko

## Abstract

The marbled crayfish (*Procambarus virginalis*) is a recently discovered freshwater crayfish species, which reproduces by apomictic parthenogenesis, resulting in a monoclonal, all-female population. The animals have become a popular source for nutritional protein in Madagascar and are increasingly being considered for commercial aquaculture. However, their potential has remained unclear and there are also significant ecological concerns about their anthropogenic distribution. We show here that the size and weight of marbled crayfish is comparable to commonly farmed freshwater crayfish. Furthermore, chemical analysis revealed a high chitin content in the marbled crayfish exoskeleton, which is a valuable source for the synthesis of chitosan and bioplastics. To allow the further evaluation of the animals in closed aquaculture systems, we developed tailored feeds that revealed an important role of methionine supplementation for animal growth. Additional analysis revealed a feed conversion rate of 1.4, which compares favorably to leading livestock for sustainable food production. Finally, we provide a concept for ecologically safe marbled crayfish aquaculture, based on key physiological characteristics that mitigate the invasive potential of the animals.

## Introduction

The marbled crayfish (*Procambarus virginalis*) is a recently discovered freshwater crayfish species that emerged in the German aquarium trade about 25 years ago^1,2^. Notably, marbled crayfish represent the only known freshwater crayfish species that reproduce by obligate apomictic parthenogenesis, a mechanism that results in the formation of an all-female, globally monoclonal population^3–5^. Through anthropogenic releases, marbled crayfish have been introduced to various freshwater systems, where they have formed numerous stable populations^6–12^.

While the introduction of marbled crayfish raises considerable ecological concerns, it also creates opportunities for human exploitation. This is exemplified by the spread of the animals on Madagascar, where their distribution area has increased 100-fold over the past 10 years^5,13,14^. This dramatic increase was largely fueled by anthropogenic distribution, as marbled crayfish have developed into a valuable source of dietary protein^15^. The rapid spread of the animals is also supported by their high tolerance in various habitat parameters and their high population densities^15^. Taken together, these characteristics suggest that marbled crayfish are an interesting candidate for aquaculture production^16^. However, their potential benefits need to be carefully balanced against their potential negative ecological impacts. As such, more data, in combination with measures that prevent the uncontrolled spread of the animals, are urgently needed.

Freshwater crayfish are increasingly popular livestock for aquaculture with a global value that now exceeds 10 billion US dollars^17^. They are a rich source of nutritional protein and contribute to the increasing global demand for it. From the more than 670 known crayfish species^18^, aquaculture production is mainly pursued with the red swamp crayfish, *Procambarus clarkii,* a species that is native to Mexico and USA^19^. In the last decade, the production of *P. clarkii* has rapidly increased^17^, as the species has proven to be robust and suited to mass production. China is currently the main producer and consumer of *P. clarkii,* with a production of more than 720,000 tons in 2015^20^.

Crayfish shells are also rich in chitin^21^. Chitin and its derivatives, such as chitosan, are important raw materials for many industries and have found frequent use in wound bandages, filter materials, and for the production of biodegradable plastics^22^. Shrimp and crab shell waste currently represents the main source for chitin production^23^.

Crayfish farming is usually done in open systems that are based on the omnivorous feeding patterns of the animals. Freshwater crayfish utilize nutrients from the bottom of the trophic food web, transfer energy to higher trophic levels, and build a major proportion of the benthos biomass^24^. The animals also mediate nutrient and energy flow of ecosystems by being prey for predators and utilizing all sources of food from the ecosystem^25–27^. Similarly, it has been described that the primary food source of marbled crayfish is autochthonous and allochthonous detritus, while other sources like zoobenthos, algae and macrophytes are utilized to a lesser extent^28^. However, the introduction of *P. clarkii* into open culture systems has created significant ecological problems, such as competition with other species and the transmission of pathogens^29^.

Nevertheless, the need for nutritional protein can create an environment that promotes commercial aquaculture of marbled crayfish. This is exemplified by the situation in Madagascar, where marbled crayfish have been deliberately released in many locations for the purpose of commercial fishery^15,30^. This development is fueled by the increasing socio-economic acceptance of marbled crayfish and their emergence as a popular food^30^. It is likely that marbled crayfish are already being considered as a livestock for commercial aquaculture in other countries and approaches that limit the ecological damage associated with marbled crayfish aquaculture are therefore urgently needed.

One important approach that limits the ecological damage of aquaculture is the use of closed systems. To facilitate aquaculture in closed systems, feeds are required that meet specific needs of the cultured animals, targeting specific production parameters (e.g., growth, feed efficiency, economics). Fish meal is often used as a preferred source of dietary protein in aquaculture, especially for new species, as it is known to be highly digestible and has a well-balanced amino acid composition. However, the use of fish meal is ecologically unfavorable. Natural alternatives, such as soybean and rapeseed meal are common alternative ingredients, but they lack methionine and lysine. Supplementation with crystalline amino acids is therefore required to meet the amino acid demands for crustaceans^31^. Our study determines key characteristics of marbled crayfish in the context of commercial aquaculture and describes tailored feeds that promote their growth in closed-system aquaculture.

## Results

### Evaluation of key characteristics for commercial aquaculture

*Procambarus clarkii* from commercial aquaculture is usually harvested at a weight of 20-25 grams^20^. We analyzed available morphometric data of marbled crayfish populations from Madagascar^15^ and additional collections from Germany. The results showed that a target weight of 20 grams is often exceeded in existing wild populations (Fig. 1A). As these populations are usually located in a challenging environment (cold climate, presence of predators, nutrient-poor habitat), their growth can likely be augmented and accelerated by aquaculture. Indeed, wild-caught animals that were grown in aquarium culture can reach weights of >50 grams (Fig. 1B). To further assess the commercial potential of marbled crayfish, we also extracted chitin from their exoskeletons. This revealed that the mean chitin content per animal of marbled crayfish shells was significantly higher than for shrimp *(Litopenaeus vannamei)* shells (2.60 % vs. 0.85 %, p<0.05, Fig. 1C). Taken together, these findings identify favorable characteristics for the commercial aquaculture of marbled crayfish.

**Figure 1.**
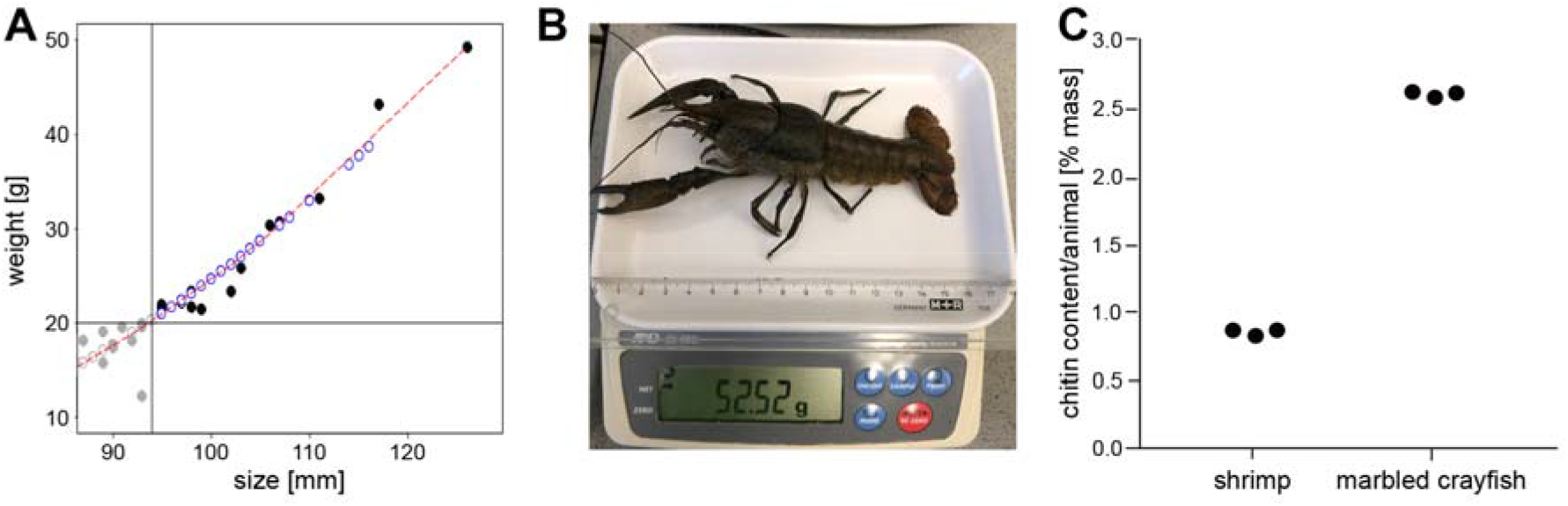
Size, weight and chitin content of marbled crayfish. (A) Morphometric data for a total of 1,537 wild-caught animals. The graph shows morphometric data for animals > 88 mm (filled circles); grey: animals <20 g; black: animals >20 g. Open circles: Size was measured, weight was estimated using regression analysis (red line) from the data where both measurements where available; grey: animals <20 g; blue: animals estimated >20 g. (B) A wild-caught and laboratory-reared specimen with a total length of 13.2 cm and a weight of 52.52 g. (C) Comparison of chitin content per animal in marbled crayfish (*P. virginalis)* shells and shrimp (*L. vannamei)* shells. An unpaired two-tailed t-test showed that the difference between the two groups is highly significant (p<0.001).

### Formulation of tailored feeds for closed-system aquaculture

As ecological concerns about marbled crayfish aquaculture remain, it is important to focus its initial development of closed systems. Closed-system aquaculture strongly benefits from the use of efficiently utilized feeds, such as tailored feeds, that are adapted to specific amino acid requirements of the animals. We therefore analyzed amino acid profiles of marbled crayfish and *L. vannamei.* In addition, the amino acid profile of a common aquarium pet feed, which was used as a control feed in this study, was also analyzed (see Methods for details). The results suggested that the control feed does not contain the ideal amino acid profile for marbled crayfish. Tailored feeds were subsequently formulated on the basis of the amino acid profiles and our experience, using a factorial modeling approach (see Methods for details). These feeds were subsequently used as a base for the test of feed supplements.

It has long been known that crayfish pigmentation is dependent on nutritionally supplied astaxanthin^32^. In a pilot experiment, we therefore determined how carotenoid supplements in our tailored feeds can affect marbled crayfish exoskeleton pigmentation. While animals that were fed with control feed appeared in darker shades of brown (Fig. 2A), the carotenoid-free tailored feed caused almost complete loss of pigmentation (Fig. 2B). The addition of astaxanthin (0.2% Carophyll Pink) resulted in a brown/blue pigmentation of the exoskeleton (Fig. 2C), which confirms the well-known effects of carotenoid supplements for our tailored feeds.

**Figure 2.**
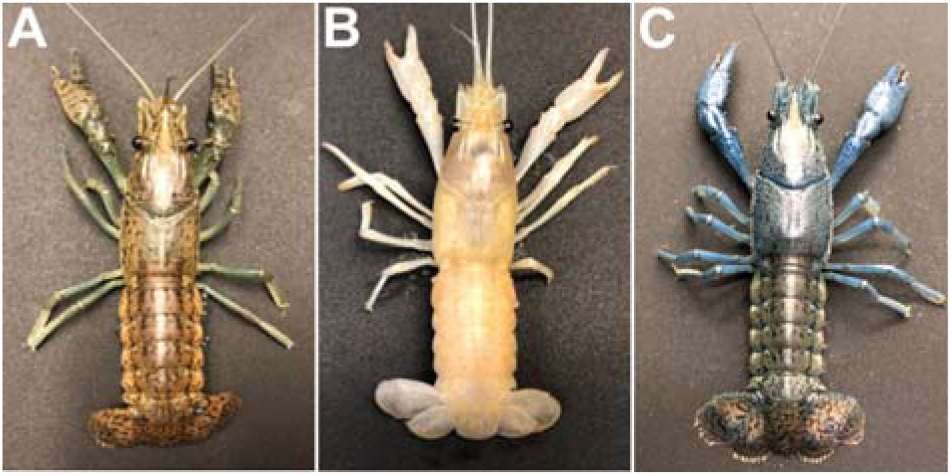
Effect of carotenoid feed supplements on the pigmentation of marbled crayfish. (A) Animal fed with control feed (D1). (B) Animal fed with a carotenoid-free tailored feed. (C) Animal fed with astaxanthin-supplemented tailored feed.

### Effects of methionine supplements on growth

Methionine is a limiting amino acid for many animals and its deficiency can directly affect animal growth^33^. To determine the optimum level methionine supplementation for marbled crayfish growth, methionine levels were varied from 0.45% to 0.7% (Tab. 1), with the control feed (D1) as a reference diet. Diet 2 (D2) is a methionine-deficient tailored feed, while D3 is a tailored feed with a methionine level that was matched to the (non-tailored) control feed. D4 was matched to the amino acid profile of the crayfish, while D5 was designed to have a methionine surplus. Our feeds were thus designed to cover specific nutritional requirements of the marbled crayfish and to determine the effects of different methionine levels on the growth of the animals.

**Table 1.**
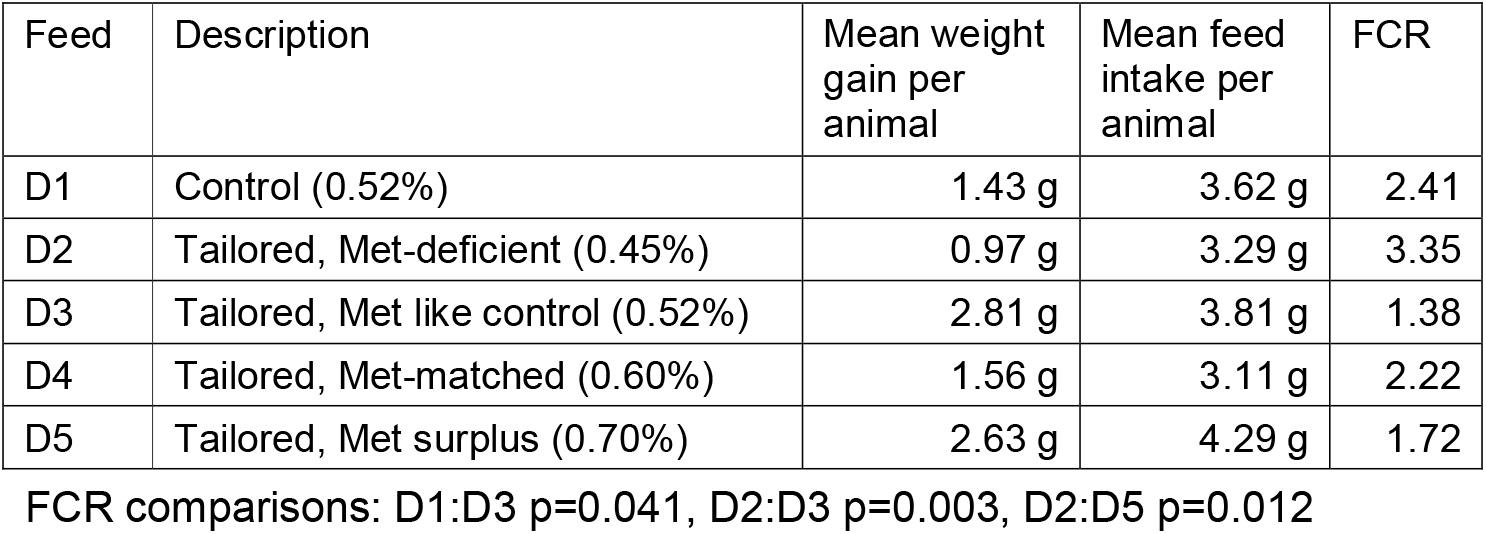
Weight gains and feed conversion ratios (FCR) of crayfish fed with different diets.

In order to test the tailored feeds and validate the methionine requirement determined in the factorial modeling approach, we designed a feeding trial, where multiple independent groups of adolescent marbled crayfish were fed for three months. No other food sources were provided to the animals. The animals were counted, measured and weighed once per week. The corrected survival probability was 52% among all groups (Fig. 3A). The low survival probability reflects the agonistic behavior of the animals, which is a known trait of freshwater crayfish, including marbled crayfish^34^. Importantly, severely injured or dead animals were removed from the trial immediately to prevent confounding effects related to cannibalistic feeding. Also, comparison between the groups (Fig. 3B) showed no significant differences, which allows the direct comparison of all groups.

**Figure 3.**
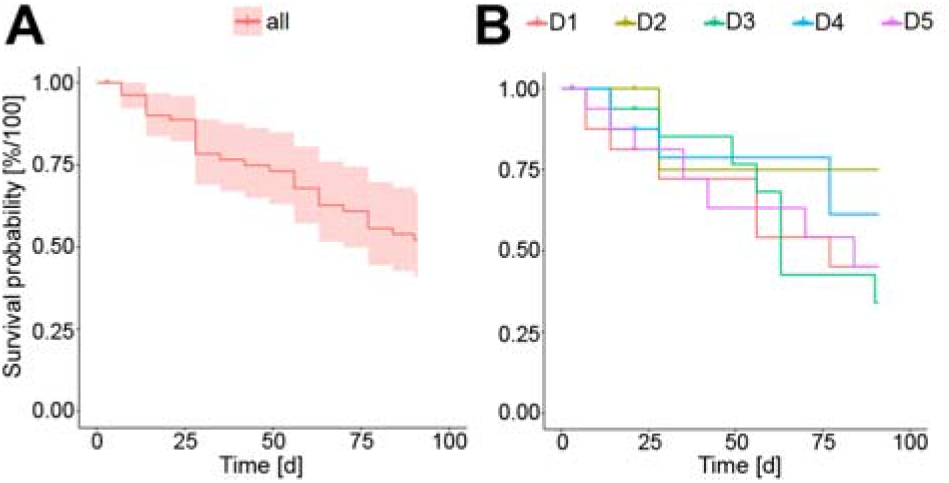
Animal survival in the feed trial. (A) Kaplan-Meier plot showing the survival probability for animals of all groups. The low overall survival probability of 52%, is due to the cannibalism of the animals, which is a known characteristic of freshwater crayfish. (B) Kaplan-Meier plot showing the survival probability for specific feeds, as indicated.

Subsequent analysis of morphometric data from the feed trial revealed a noticeable effect of the tailored feeds on the growth of the animals. When compared to the non-tailored control feed (D1), feed D3 (tailored, methionine matched) resulted in an increased weight gain (Fig. 4). A comparison of all tailored feeds showed that animals fed with D5 (surplus methionine) and D3 (methionine matched to control) showed a faster weight gain compared to D2 (methionine deficient) and D4 (methionine matched to requirement). During the early stages of the trial (until week 8), D5 showed the best performance, while D3 showed the best performance during the late stages of the trial (Fig. 4).

**Figure 4.**
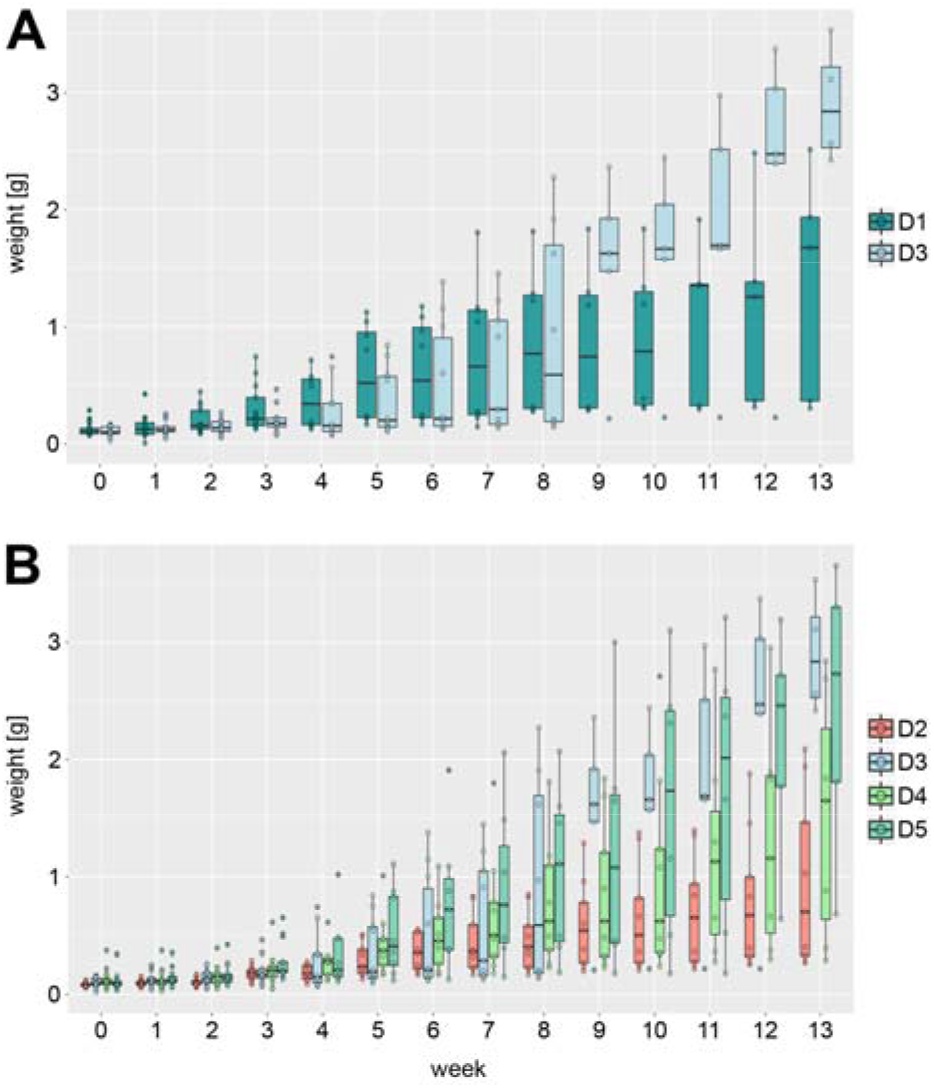
Effect of methionine feed supplements on the growth of marbled crayfish. Box plots showing the weight per group and week. (A) Comparison of the control feed (D1) with the matched tailored feed (D3). The tailored feed D3 shows a higher weight gain compared to D1. (B) Comparison of all tailored feeds (D2-D5). D2 shows the lowest weight gain over time. D3 and D5 have the highest weight gain, but for D5 the weight gain is stronger at the beginning of the trial until week 9. From week 9 till the end of the trial D3 shows the highest weight gain. Both groups have similar weight gains after 13 weeks with 2.81 g/animal for D3 and 2.63 g/animal for D5.

### Determination of feed conversion ratios

At the end of the trial, a mean weight gain of 2.81 g (min: 2.3 g; max: 3.45 g; SD: 0.46) was observed for D3, while a mean gain of only 1.43 g (min:0.89 g; max: 2.38 g; SD: 0.68) was observed for D1 (Fig. 4). Consistently, the total weight gain of D3 was more than double weight gain achieved with D1 (Tab. 1). Subsequent determination of feed conversion ratios (FCR) revealed that the FCR for D3 was significantly (p<0.05. ANOVA) better than for D1 (1.38 to 2.41, Tab. 1). These results again illustrate the superior performance of tailored feeds. Also, D2 (methionine-deficient) showed the lowest FCR among all feeds (Tab. 1), which again illustrates the importance of methionine supplementation for marbled crayfish feeds. The calculated FCR for D3 (1.38, Tab. 1) compares very well to other arthropod species that are being considered as leading candidates for the sustainable production of nutritional protein^35^.

## Discussion

The increasing demand for nutritional protein and bioplastics requires the development of sustainable and ecologically friendly production strategies. Our size and weight data of marbled crayfish revealed comparable features to commercially harvested *P. clarkii,* which is the dominant stock in crayfish aquaculture^20^, consistent with the close genetic relationship of the two species^36^. Additional potential for commercialization was provided by the observation that marbled crayfish shells contain 3 times more chitin per animal compared to *L. vannamei* shells, which currently represent the main source of commercially produced chitin. In addition, our results establish marbled crayfish as highly efficient feed converters, which further illustrates their potential for sustainable aquaculture production.

However, important questions about the ecological impact of marbled crayfish remain to be resolved. In particular, the further release of the species should be prevented to mitigate their potential invasiveness. To allow marbled crayfish aquaculture in closed systems, tailored feeds were formulated. When diets are imbalanced and do not meet the animal’s requirements for all amino acids, part of the amino acids are catabolized for energy rather than for protein synthesis^37,38^. This leads to inadequate protein and feed utilization and high ammonia production in the water^37,38^. Our findings suggest that tailored feeds, optimized to the amino acid profile of the marbled crayfish, and with adjusted methionine concentrations, can improve the growth performance of cultured marbled crayfish considerably.

Methionine is often the limiting amino acid to promote growth in aquatic animals if feeds contain mainly plant proteins^33^. To cover these needs, different feeds with various methionine levels were tested. Indeed, the lowest growth rate was detected for the feed with the lowest methionine level. This shows that methionine is a growth-limiting nutrient for marbled crayfish, similar to other crayfish, shrimps and fish^31,39,40^. Interestingly, the feed with the highest methionine level mainly promoted growth at earlier developmental stages, while later stages performed best with a feed that had an intermediate methionine level. These observations indicate different needs of methionine in different stages of crayfish development. Studies in other aquatic animals show that growth rates reach a plateau^41,42^, or even decrease when the methionine requirements of the animals are overfed^43,44^. Our results suggest that methionine levels of 0.7% optimally support the growth of juvenile and early adolescent marbled crayfish. A balanced methionine supplementation is particularly relevant in a hatchery and nursery environment, where it supports rapid growth while maintaining high water quality.

The use of closed aquaculture systems is important to prevent the release of marbled crayfish into their environment (Fig. 5). While this is feasible and cost-effective for the hatchery and nursery stages, closed systems would represent a major economic disadvantage for the growout stage, which requires large surfaces. However, marbled crayfish are known to have very low salinity tolerance^45^. Based on the available information^45^, salt concentrations that are commonly used for shrimp (*L. vannamei)* aquaculture, would effectively prevent the spread and reproduction of marbled crayfish. The placement of marbled crayfish growout ponds in brackish water environments (Fig. 5) should therefore effectively mitigate their invasive spread. Further safeguarding and environmental monitoring of marbled crayfish aquaculture can be achieved by the use of established detection methods that are based on environmental DNA^46^.

**Figure 5.**
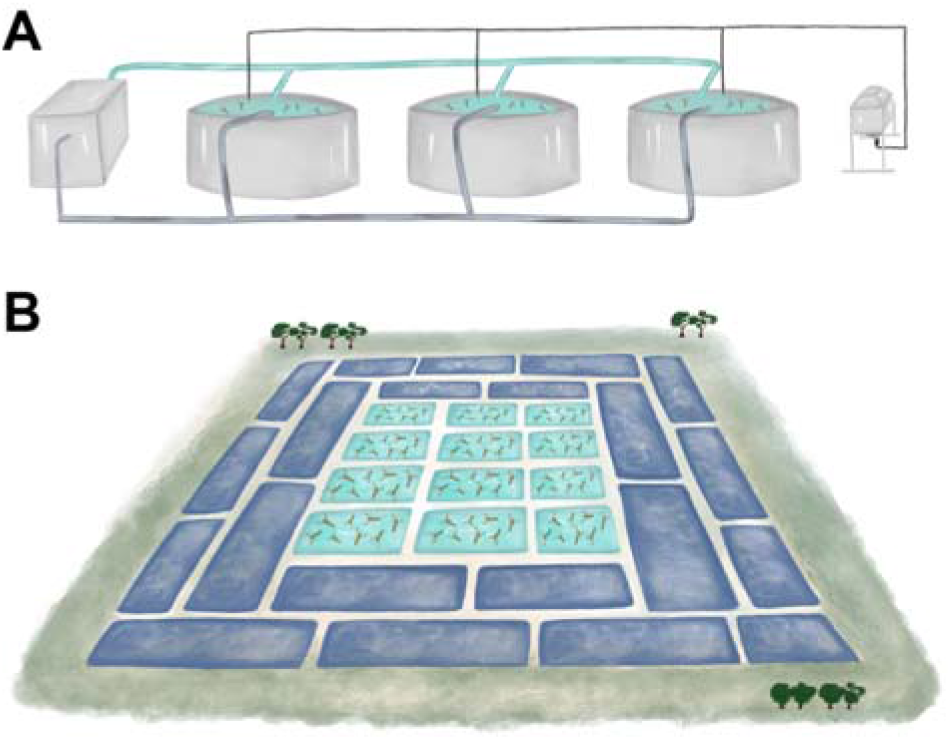
Schematic illustration of closed-system marbled crayfish aquaculture. (A) A recirculating aquaculture system for the hatchery/nursery of marbled crayfish juveniles and adolescents. The box on the left side illustrates the filter system, the container on the right side represents the feeding system. (B) Freshwater growout ponds (turquoise) for marbled crayfish aquaculture, located in a brackish water (dark blue) aquaculture environment. The low salinity tolerance of marbled crayfish^45^ effectively closes the system.

## Methods

### Morphometric data analysis

Morphometric data of various populations in Madagascar^15^ and Germany (Reilinger See, 49.296893N, 8.544591 E; Murner See, 49.348875 N, 12.206505E) was analyzed. To predict marbled crayfish growth rates, a local polynomial regression using the locally estimated scatterplot smoothing (LOESS) method in R (version 3.6.1) was performed on the total length and weight of the animals (n=1,537). The prediction was performed by using the predict function on the total length of the animals (n=347) from Reilinger See (49.296893N, 8.544591E).

### Isolation of chitin

The following procedure is based on slightly modified literature protocols^47,48^ to remove CaCO_3_ and protein from shellfish shells. Peeled exoskeletons from pre-cooked marbled crayfish and shrimps (*L. vannamei)* were collected and air dried. Exoskeletons were subsequently ground to powder with a Thermomix^®^ (level 8, 30 s). From each species, 7 g of powdered exoskeletons were weighed out and added portion wise to a magnetically stirred solution of 1 M HCl (50 ml) in a wide necked 250 ml Erlenmeyer flask at 23 °C. Note: care must be taken to add the material slowly as vigorous foaming occurs, which can cause overflowing. After the addition was complete, an additional 10 ml of 1 M HCl was used to rinse the sides of the flask to ensure all material was submerged. After 75 min, the resulting slurry was vacuum filtered through a porosity #3 sintered glass Büchner funnel using a membrane pump as a source of vacuum. The solid was repeatedly rinsed with deionized H_2_O (6 x 50 ml) until the filtrate was neutral to pH indicator paper. After air drying with vacuum on the filter for 15 min, the powder was transferred to a 250 mL Erlenmeyer flask containing 1 M NaOH (75 mL), where it was magnetically stirred at 23 °C. After 24 h, the resulting slurry was vacuum filtered through a porosity #3 sintered glass filter Büchner funnel as before. The solid was repeatedly rinsed with deionized H_2_O (6 x 50 mL) until the filtrate was neutral to pH indicator paper. The resulting solid was air dried with vacuum for ~20 min, transferred to a 100 mL round bottom flask, dried under high vacuum (~5 x 10^−2^ mbar) for 24 h, and weighed on an analytical balance. This protocol was performed in triplicate for each species.

### Determination of amino acid requirements using factorial modeling

One randomly selected crayfish (14.77 g), which had been fed standard crayfish pet feed was decapitated and freeze-dried (final mass: 4.35 g). The whole-body amino acid profile of this sample (Tab. S1) was analyzed using ion-exchange chromatography (AMINOLab^®^, Evonik Nutrition & Care, Germany), except for tryptophan, which was estimated using high-performance liquid chromatography (HPLC). In addition, the amino acid content of a common aquarium fish feed (NovoPleco, JBL, Neuhofen, Germany), which was used as a control feed, was also analyzed (Fig. S1). Using the amino acid profile of the crayfish, the amino acid composition was calculated for a body weight gain of 1.5 g. The utilization of different amino acids absorbed across the gut was assumed to be 50-60%, and the maintenance requirements for different amino acids were considered to be 15-25% of the amino acids absorbed (i.e., on digestible basis). Amino acid requirements calculated with a factorial modeling approach are presented in Tab. 1 on digestible basis and total basis (assuming 85% digestibility for amino acids), following established procedures. Absolute requirement values derived as mg were converted into % feed assuming 1.5 g as the feed requirement for 1.5 g body weight gain.

### Experimental diets

The detailed ingredient and nutrient composition of the diets used in this study are provided in Tab. S2. A basal diet was formulated using soybean meal, soy protein concentrate, fish meal and krill meal as the main protein sources. A basal diet (D2) was formulated to contain 29% crude protein and 18.31 MJ/kg gross energy. The amino acid profile of the diet except for methionine was balanced considering the requirements predicted using factorial modeling approach (Table 1). The basal diet (D2) was formulated to be low in Met (0.45%) and Met + Cys (0.86%) and was supplemented with increasing levels of methionine dipeptide (AQUAVI^®^ Met-Met): 0.07% (D3), 0.15% (D4), and 0.25% (D5). Feed contents were validated by amino acid analysis (Tab. S2). The D2-D5 feeds were produced in pellets of 2 mm size.

### Feed trial

Each feed was tested for three months. In total, 100 adolescent animals (mean total length: 1.75 cm, SD: 0.25 cm; mean weight: 0.11 g, SD: 0.05 g) from our laboratory stock^49^ were used in the trial. Per feed, 20 animals were kept in 4 tanks (25.6 x 18.1 x 13.6 cm), with five animals per tank, at 20 °C under natural daylight. The animals were fed daily at 17:00 with 0.08 g of feed. Higher amounts of feed were offered occasionally but refused by the animals. Because of the specific feeding behavior of crayfish (prolonged feeding time; preference to stay hidden), it was not possible to determine accurate feed intake. A fixed amount of feed was provided in each tank, and no uneaten feed was recovered. Daily mean feed intake per tank was calculated by dividing the amount of feed fed to a tank on a given day by the number of animals survived on that day. Calculated daily mean feed intake value was summed over the whole experimental period to calculate the total mean feed intake in each tank. All animals were measured and weighed once per week. Water parameters were checked daily for temperature and once per week for NH_4_, NO_3_, NO_2_ and O_2_ (JBL Test sets, Neuhofen, Germany). The results confirmed good water quality for the entire duration of the trial (Tab. S3).

### Feed trial data analaysis

Survival probabilities for all animals and the groups D1 to D5 were calculated in R using the CRAN packages survival (version 2.44-1.1) and survminer (version 0.4.6). Kaplan-Meier plots were generated, and survival probabilities were calculated for each feed group and for all animals. To investigate differences in survival between feeds pairwise p-values were calculated using a log-rank test. Finally, p-values were adjusted using the Benjamini-Hochberg procedure. To assess differences in the means of feed conversion rates among the five feed groups a one-way analysis of variances (ANOVA) was performed. As the total variances between groups were statistically significant (p-value = 0.00239) a more detailed pairwise comparison between all groups was performed using Tukey’s HSD test.

## Acknowledgements

We thank Ranja Andriantsoa and Frank Lenich for morphometric data.

## Author contributions

S.T. performed the experiments and analyzed the data. K.M. designed the feeds and the feed trial and analyzed the data. J.G. performed statistical analyses and growth predictions. J.L. and A.K.M. performed the chitin extraction. F.B. and F.L. conceived the study. S.T. and F.L. wrote the paper with input from the other authors. All authors read and approved the final manuscript.

