## Supplementary Information for "Physiological properties and tailored feeds to support aquaculture of marbled crayfish in closed systems"

Sina Tönges, Karthik Masagounder, Julian Gutekunst, Jasmin Lohbeck, Aubry K. Miller, Florian Böhl, and Frank Lyko

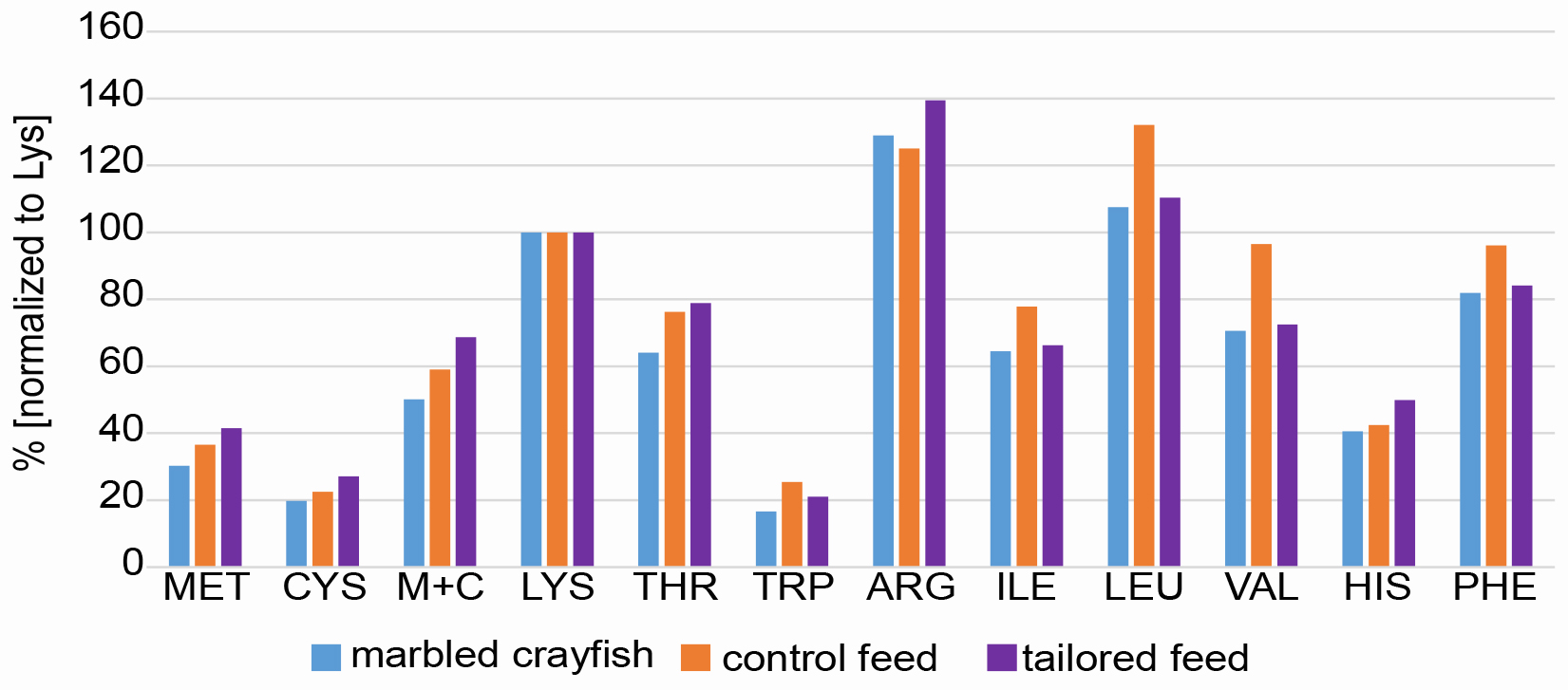

Figure S1. Comparison of amino acid profiles for the development of tailored feeds. Bars show relative amino acid levels for marbled crayfish (blue), control feed (orange) and tailored feeds (purple). All profiles are normalized to lysine (100%).

Table S1. Amino acid profiles of shrimp (*Litopenaeus vannamei*) and marbled crayfish (*Procambarus virginalis*).

| **Shrimp (N=315)** | **CP** | **Met** | **Cys** | **MC** | **Lys** | **Thr** | **Trp** | **Arg** | **Ile** | **Leu** | **Val** | **His** | **Phe** |
| --- | --- | --- | --- | --- | --- | --- | --- | --- | --- | --- | --- | --- | --- |
| Dry matter [%] | 66.51 | 1.42 | 0.58 | 2.01 | 4.39 | 2.28 | 0.62 | 5.05 | 2.49 | 4.29 | 2.80 | 1.32 | 3.23 |
| CP [%] | 100 | 2.14 | 0.88 | 3.02 | 6.61 | 3.43 | 0.93 | 7.59 | 3.75 | 6.45 | 4.20 | 1.99 | 4.85 |
| IAA [%] |  | 32 | 13 | 46 | 100 | 52 | 14 | 115 | 57 | 98 | 64 | 30 | 73 |
| **Crayfish (N=1)** | **CP** | **Met** | **Cys** | **MC** | **Lys** | **Thr** | **Trp** | **Arg** | **Ile** | **Leu** | **Val** | **His** | **Phe** |
| Dry matter [%] | 41.97 | 0.75 | 0.49 | 1.24 | 2.48 | 1.59 | 0.41 | 3.20 | 1.60 | 2.66 | 1.75 | 1.00 | 2.03 |
| CP [%] | 100 | 1.79 | 1.17 | 2.96 | 5.91 | 3.78 | 0.98 | 7.62 | 3.81 | 6.35 | 4.17 | 2.39 | 4.84 |
| IAA [%] |  | 30 | 20 | 50 | 100 | 64 | 17 | 129 | 65 | 108 | 71 | 41 | 82 |
|  |  | -2 | 7 | 4 | - | 12 | 3 | 14 | 8 | 10 | 7 | 11 | 9 |

MC: Methyionine + Cysteine

Table S2. Ingredient and nutrient composition of the diets used in this study.

| **Ingredients (%)** | **D1** | **D2** | **D3** | **D4** | **D5** |
| --- | --- | --- | --- | --- | --- |
| Wheat flour | n.a. | 45.21 | 45.21 | 45.21 | 45.21 |
| Soybean meal 45%CP | n.a. | 25.59 | 25.59 | 25.59 | 25.59 |
| Soy protein concentrate | n.a. | 8.14 | 8.14 | 8.14 | 8.14 |
| Wheat bran | n.a. | 8.12 | 8.12 | 8.12 | 8.12 |
| Fish meal 60%CP | n.a. | 5.00 | 5.00 | 5.00 | 5.00 |
| Soy lecithin | n.a. | 2.23 | 2.23 | 2.23 | 2.23 |
| Krill meal | n.a. | 2.00 | 2.00 | 2.00 | 2.00 |
| Fish oil | n.a. | 1.00 | 1.00 | 1.00 | 1.00 |
| Monocalciumphosphate | n.a. | 0.71 | 0.71 | 0.71 | 0.71 |
| Vitamin-Mineral Premix | n.a. | 1.00 | 1.00 | 1.00 | 1.00 |
| Astaxanthin | n.a. | 0.20 | 0.20 | 0.20 | 0.20 |
| Aquavi® Met-Met | n.a. | 0.00 | 0.07 | 0.15 | 0.25 |
| Biolys® (L-Lys Sulphate) | n.a. | 0.12 | 0.12 | 0.12 | 0.12 |
| ThreAMINO® (L-Thr) | n.a. | 0.24 | 0.24 | 0.24 | 0.24 |
| L-Arginine | n.a. | 0.36 | 0.36 | 0.36 | 0.36 |
| L-Histidine | n.a. | 0.08 | 0.08 | 0.08 | 0.08 |
| Dry matter | n.a. | 89.56 (92.51) | 89.56 (91.57) | 89.56 (91.87) | 89.56 (90.25) |
| Crude protein | (29.00) | 29.37 (29.32) | 29.37 (29.46) | 29.37 (30.47) | 29.37 (29.77) |
| Crude fat | n.a. | 6.46 | 6.46 | 6.46 | 6.46 |
| GE (MJ/kg) | n.a. | 18.31 | 18.31 | 18.31 | 18.31 |
| Lys | (1.43) | 1.60 (1.46) | 1.60 (1.48) | 1.60 (1.56) | 1.60 (1.53) |
| Met | (0.52) | 0.45 (0.43) | 0.52 (0.50) | 0.60 (0.62) | 0.70 (0.68) |
| Cys | (0.32) | 0.42 (0.41) | 0.42 (0.41) | 0.42 (0.42) | 0.42 (0.41) |
| Met+Cys | (0.84) | 0.86 (0.84) | 0.91 (0.91) | 0.96 (1.04) | 1.01 (1.09) |
| Thr | (1.09) | 1.27 (1.22) | 1.27 (1.22) | 1.27 (1.25) | 1.27 (1.22) |
| Trp | (0.36) | 0.37 (n.a.) | 0.37 (n.a.) | 0.37 (n.a.) | 0.37 (n.a.) |
| Arg | (1.79) | 2.24 (2.10) | 2.24 (2.11) | 2.24 (2.16) | 2.24 (2.11) |
| Ile | (1.11) | 1.19 (1.15) | 1.19 (1.17) | 1.19 (1.21) | 1.19 (1.20) |
| Leu | (1.89) | 2.06 (1.98) | 2.06 (2.00) | 2.06 (2.07) | 2.06 (2.03) |
| Val | (1.38) | 1.32 (1.27) | 1.32 (1.29) | 1.32 (1.33) | 1.32 (1.31) |
| His | (0.61) | 0.80 (0.75) | 0.80 (0.76) | 0.80 (0.79) | 0.80 (0.76) |
| Phe | (1.37) | 1.35 (1.33) | 1.35 (1.33) | 1.35 (1.37) | 1.35 (1.35) |

Analyzed dry matter, crude protein and amino acid content of feeds are provided in parentheses. n.a.: not analyzed. Amino acids are indicated in standard three-letter code.

Table S3. Water parameters during the feed trial.

| **Parameter** | **Results** |
| --- | --- |
| Temperature [°C] | 20 ±1 |
| O_2_ [mg/l] | 8 for all measurements |
| NH_4_ [mg/l] | <0.2 for all measurements |
| NO_3_ [mg/l] | <1 for all measurements |
| NO_2_ [mg/l] | <0.1 for all measurements |
